# Natural variation in *PHY* causes differential response to R/FR light following a latitudinal cline in Norway spruce

**DOI:** 10.1101/2022.08.28.505569

**Authors:** Sonali Sachin Ranade, María Rosario García-Gil

**Affiliations:** Department of Forest Genetics and Plant Physiology, Umeå Plant Science Centre, Swedish University of Agricultural Sciences, SE-901 83 Umeå, Sweden

**Keywords:** cline, cryptochrome, light quality, local adaptation, missense mutation, Norway spruce, photoreceptor, phytochrome, polymorphism, SNP

## Abstract

Detection of the genomic basis of local adaptation to environmental conditions is challenging in forest trees. Phytochromes (*PHY*) and cryptochromes (*CRY*) perceive the red (R)/far-red (FR) and blue light respectively and, play a fundamental role in regulating the light pathway in plants contributing to plant growth and development. Norway spruce shows an adaptive latitudinal cline for shade (low R:FR or FR-enriched light) tolerance and requirement of FR light for its growth, thus showing differential response to light quality. We analyzed the exome capture data that included a uniquely large data set of 1654 Norway spruce trees sampled from different latitudes across Sweden that differ in exposure to photoperiod and FR light received during the growing season. Statistically significant clinal variation was detected in allele and genotype frequencies of missense mutations in coding regions belonging to well-defined functional domains of *PHYO* (PAS-B), *PHYP2* (PAS fold-2), *CRY1* (CCT1) and *CRY2* (CCT2) that strongly correlates with the latitudinal gradient in response to variable light quality in Norway spruce. Asn835Ser in *PHYO* displayed the steepest cline among all other polymorphisms. We propose that these variations represent signs of local adaptation to light quality in Norway spruce.

## Introduction

Light plays a vital role in the regulation of plant growth and development. Phytochrome (*PHY*) and cryptochrome (*CRY*) code for the key photoreceptors that perceive the red (R)/far-red (FR) and blue light respectively and, play fundamental roles in regulating the developmental processes throughout the plant life cycle by sensing the light quality, photoperiod and modulating the light signaling pathway (Casal 2000). *PHY* and *CRY* mutants have been described in plant model systems e.g. *Arabidopsis thaliana* (*Arabidopsis*) that are sensitive to R/FR wavelength (Franklin and Quail 2010; Whitelam and Devlin 1997) and blue light (Xuhong Yu 2010). Unlike the angiosperm model plant *Arabidopsis*, conifer *PHY*s and *CRY*s are not well characterized for the molecular mechanisms regarding their mode of action regulating different light sensing pathways. Five *PHY*s have been characterized in *Arabidopsis* – *PHYA, PHYB, PHYC, PHYD* and *PHYE* (Casal 2000). There are only two major *PHY*s denoted as *PHYO* and *PHYP* in conifers, while *PHYN* is a subtype of *PHYO* (Mathews 2010). *PHYO* phylogenetically diverged leading to *PHYA* and *PHYC* in angiosperms; likewise, *PHYP* phylogenetically diverged leading to *PHYB, PHYD* and *PHYE* in angiosperms (Schneider-Poetsch et al. 1998). Similarly, the three *CRY*s are well studied in *Arabidopsis* – *CRY1, CRY2* and *CRY3* (Ponnu and Hoecker 2022), whereas in conifers only *CRY1* and *CRY2* have been reported so far (Pashkovskiy et al. 2021; Ranade et al. 2019a; Ranade and García-Gil 2013).

The R:FR ratio under sunlight is 1.2 (Warrington et al. 1989; Smith 1994). Under the vegetative shade, there is a decrease in the R:FR ratio (0.2–0.8) as R is absorbed by the chlorophyll and other leaf pigments, whereas the FR is reflected (Ballare et al. 1987). Plants perceive shade as a decrease in the R:FR ratio, where there is a higher FR than R. Species that become established and compete well in fully shaded conditions are termed shade tolerant (Grebner et al. 2021). Shade conditions are similar to astronomic shade or twilight, which is also characterized by a low R:FR ratio (Nilsen 1985) or the FR-enriched light. The geographic location of Sweden leads to a pronounced latitudinal difference in the duration of twilight when the northern latitudes receive longer daily exposure to FR-enriched light (twilight) as compared to the southern latitudes during the growing season (Supplementary Information, Figure S1 and references therein).

Local adaption in plants tends to render higher mean fitness to the local populations in their native environment. Detection of the genomic basis of local adaptation to environmental conditions is challenging in forest trees especially in the long-lived and ancient conifers with large genome sizes, such as Norway spruce (*Picea abies* [L.] H. Harst). Norway spruce is one of the most important conifer species of the Boreal forest that has high economical value. Norway spruce is shade tolerant and it shows a requirement for far-red light to maintain the growth that follows a latitudinal gradient (Clapham et al. 1998; Mølmann et al. 2006); however, the underlying mechanism or any genetic correlation with the phenomenon has not been demonstrated. It is also evident from our previous study that Norway spruce shows the presence of an adaptive cline for shade or FR-enriched light (low R:FR ratio) (Ranade and García-Gil 2021). Norway spruce seedlings from latitudes across Sweden show significant variation in hypocotyl length when grown under different R:FR ratios (Supplementary Information, Figure S2 and references therein). Norway spruce is shade tolerant but neither *PHY*s nor *CRY*s were found to be involved in the shade tolerance in the species (Ranade et al. 2019b). In addition, neither *PHY*s nor *CRY*s were detected to be differentially regulated under shade in the two populations from different latitudes in Norway spruce (Ranade and García-Gil 2021). Recently, enhanced lignin synthesis and ecotypic variation in defense-related gene expression were reported in Norway spruce in response to shade; the study discussed the local adaptation to extended FR-enriched light as a potential reason for the differential defense-related gene expression (Ranade et al. 2022). Photoreceptors perceive the light quality and they are the key elements of the light pathway. Therefore, we were interested in analyzing the SNPs present in the photoreceptors and to determine if those SNPs follow a latitudinal gradient in Norway spruce that correlates well with the adaptive cline to variable light conditions across populations present in various latitudes that are exposed to differential light quality due to their geographic location.

With the availability of the complete genome of Norway spruce (Nystedt et al. 2013), we revisited the classification of the *PHY*s and *CRY*s in this species and analyzed their phylogeny in this work.

## Material and methods

### PHYs and CRYs from Norway spruce

Norway spruce homologs for the PHYs and CRYs were retrieved from PlantGenIE (https://plantgenie.org) by performing Blastp with the corresponding *Arabidopsis* members from TAIR (https://www.arabidopsis.org/index.jsp). PHYs and CRYs from other species were retrieved from PlantGenIE (https://plantgenie.org) and NCBI (https://www.ncbi.nlm.nih.gov/). The details of all the sequences included in this study are presented in Table S1 (Supplementary Information). The domain regions of PHYs and CRYs from Norway spruce (Supplementary Information, Figures S3-S11) were confirmed by referring to its best match in *Arabidopsis* and by performing searches in the Conserved Domain Database (CDD) (Marchler-Bauer et al. 2015) and UniProt (Bateman et al. 2021), referring to the literature (Li et al. 2011; He et al. 2015; Klar et al. 2007) and by aligning them with *Arabidopsis* sequences using MUSCLE (Edgar 2004). Phylogenetic trees of protein sequences of PHYs and CRYs from Norway spruce along with the PHYs and CRYs from a few model plants (e.g. *Arabidopsis*, soybean, eucalyptus and poplar) were constructed for further validation, using phylogeny.fr in the ‘one click mode’ using default settings (https://www.PHYlogeny.fr/) (Dereeper et al. 2008). In brief, the alignment was done with MUSCLE (Edgar 2004), phylogeny was done using PHYML (Guindon et al. 2010) which is based on the maximum-likelihood principle and the phylogenetic tree was prepared using TreeDyn (Chevenet et al. 2006).

### Plant material, DNA extraction and sequencing

A total of 1654 individuals (unrelated parents) originating from different latitudes across Sweden, were included in this study for analysis of the SNP variation following a latitudinal cline (Ranade and García-Gil 2021). The 1654 individuals belong to progeny trials established from the half-sib families from wild trees growing at the same latitudes as the plantations, which implies that these plantations are not domesticated trees, and therefore they reflect the natural variation. Only one tree per progeny was sampled and the mothers of each progeny were unrelated. For the current analysis, trees were divided into six populations, S1-S6. S1 comprised 245 trees from latitudes 55-57, S2 – 213 trees from latitude 58, S3 – 187 trees from latitudes 59-60, S4 – 213 trees from latitudes 61-62, S5 – 573 trees from latitudes 63-64 and S6 – 223 trees from latitudes 65-67. This categorization of the population was done taking into consideration the fact that the quality of light differs latitude-wise from south to north in Sweden; during the growing season (summer), the northern latitudes receive an extended period of FR light as compared to the southern ones.

Details regarding DNA extraction, exome capture, genotyping and SNP annotation have been previously described in Baison et al. (Baison et al. 2019). In short, total genomic DNA was extracted from 1654 trees, from dormant buds, using the Qiagen Plant DNA extraction (Qiagen, Hilden, Germany) following the manufacturer’s protocol and DNA quantification was performed using the Qubit ds DNA Broad Range (BR) Assay Kit (Eugene, Oregon, USA). DNA library preparation and exome capture sequencing were performed at the RAPiD Genomics (Gainesville, Florida, USA). Sequence capture was performed using the 40, 018 diploid probes designed and evaluated for *P. abies* (Vidalis et al. 2018). Sequencing was performed using an Illumina HiSeq 2500 instrument (San Diego, California, USA) on the 2 × 100 bp sequencing mode.

### Determination of nucleotide variation in PHYs and CRYs

Raw reads were mapped against the Norway spruce reference genome v.1.0 and variant calling was performed using GATK HAPLOTYPECALLER v.3.6 (Van der Auwera et al. 2013). The resultant SNPs were annotated using default parameters for SNPEFF 4 (Cingolani et al. 2012). The vcf file was filtered using settings; --min-alleles 2 --max-alleles 2 --maf 0.01 --remove-indels --minQ 10 --max-missing 0.9. *PHYO, PHYN, PHYP1, PHYP2, CRY1, CRY2* and *CRY3* were analyzed for the presence of SNPs and possible clines associated with those SNPs. Only bi-allelic SNPs were included in this study. The vcf file containing data from the exome sequencing results of 1654 trees that includes the photoreceptors from the current analysis, is deposited in Zenodo, which is the open-access repository developed under the European OpenAIRE program and operated by CERN. Allele frequencies, genotype frequencies and Hardy-Weinberg equilibrium (HWE) *P-values* were determined using SNPassoc statistical package (Gonzalez et al. 2007). ANOVA and Tukey’s *post-hoc* tests (Bonferroni *P-values*) were applied to determine the statistical significance of the difference in the allele and genotype frequencies across the populations included in the study. Genetic diversity among the six different populations (pairwise *F*_*ST*_ estimates) was estimated using DnaSP 6 (Rozas et al. 2017) including both the synonymous + missense SNPs detected within *PHYO, PHYP2, CRY1* and *CRY2*. Allele frequencies in each population regarding these four genes were calculated and then regressed on population latitude. R^2^ of the linear regression was computed as the proportion of total variance of latitude explained by the frequency of each marker (Berry and Kreitman 1993). R^2^ is the goodness-of-fit of the linear regression model.

## Results

### Phylogeny of PHYs and CRYs

The phylogenetic analysis is in accordance with the previous work in this area (Mathews 2010) and it also gives additional information. The phylogenetic tree of phytochromes (Figure 1) shows that there are three PHYs (PHYP, PHYN and PHYO) in Norway spruce as described earlier. It shows two PHYPs (PHYP1 and PHYP2) in Norway spruce similar to poplar where two PHYBs (PHYB1 and PHYB2) were identified (Howe et al. 1998). Cryptochromes have not been described extensively earlier. The CRY phylogeny (Figure 2) shows that Norway spruce has CRY1, CRY2 and two CRY3 genes. Details of the sequences included in the trees are included in the supplementary information (Table S1). The Figures S3-S11 in the supplementary information represent the alignments of Norway spruce PHYs and CRYs with the respective members in *Arabidopsis*, which shows that the photoreceptors are well conserved. Full-length sequences were detected for all the photoreceptors except for both the CRY3 sequences that were found to be partial. In the case of PHYO, PlantGenIE searches retrieved three partial sequences - MA_6809p0010, MA_6809p0020, and MA_6809p0030, which were annotated as PHYA. However, these three sequences were combined to form the full-length PHYO protein, which is well conserved with *Arabidopsis* PHYA (Supplementary Information, Figure S4). This combined sequence is denoted as MA_6809 in the current work.

**Figure 1.**
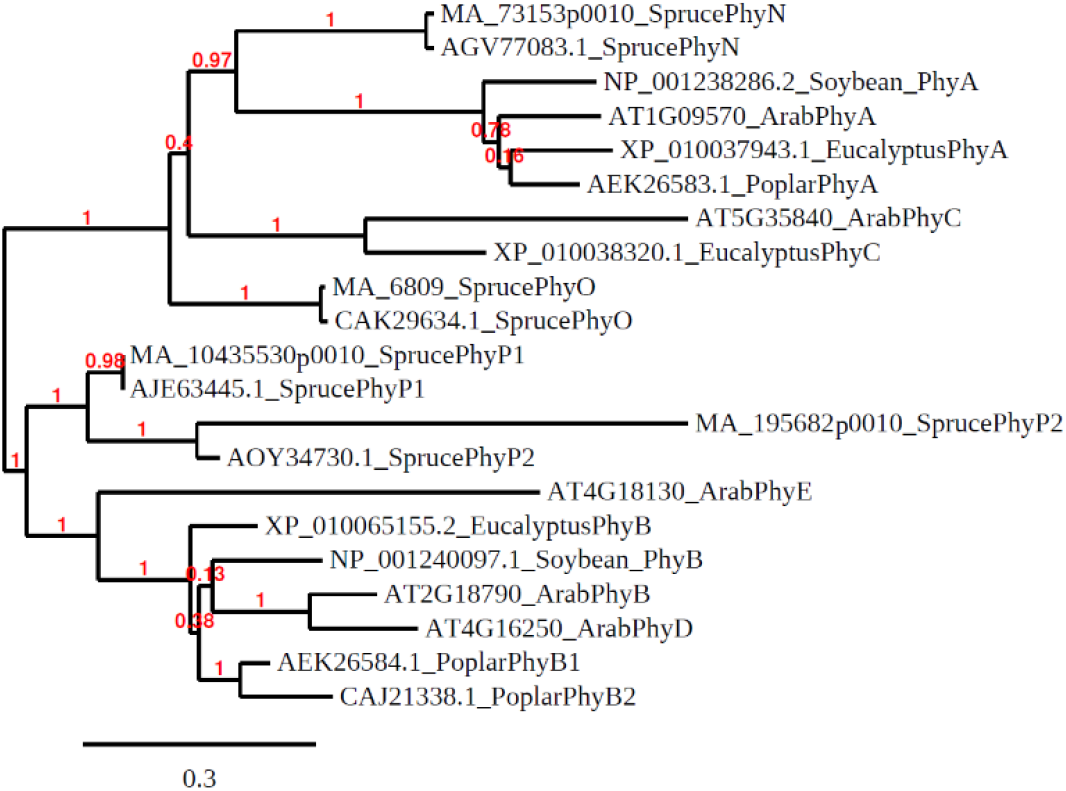
Phylogenetic tree of phytochromes in Norway spruce

**Figure 2.**
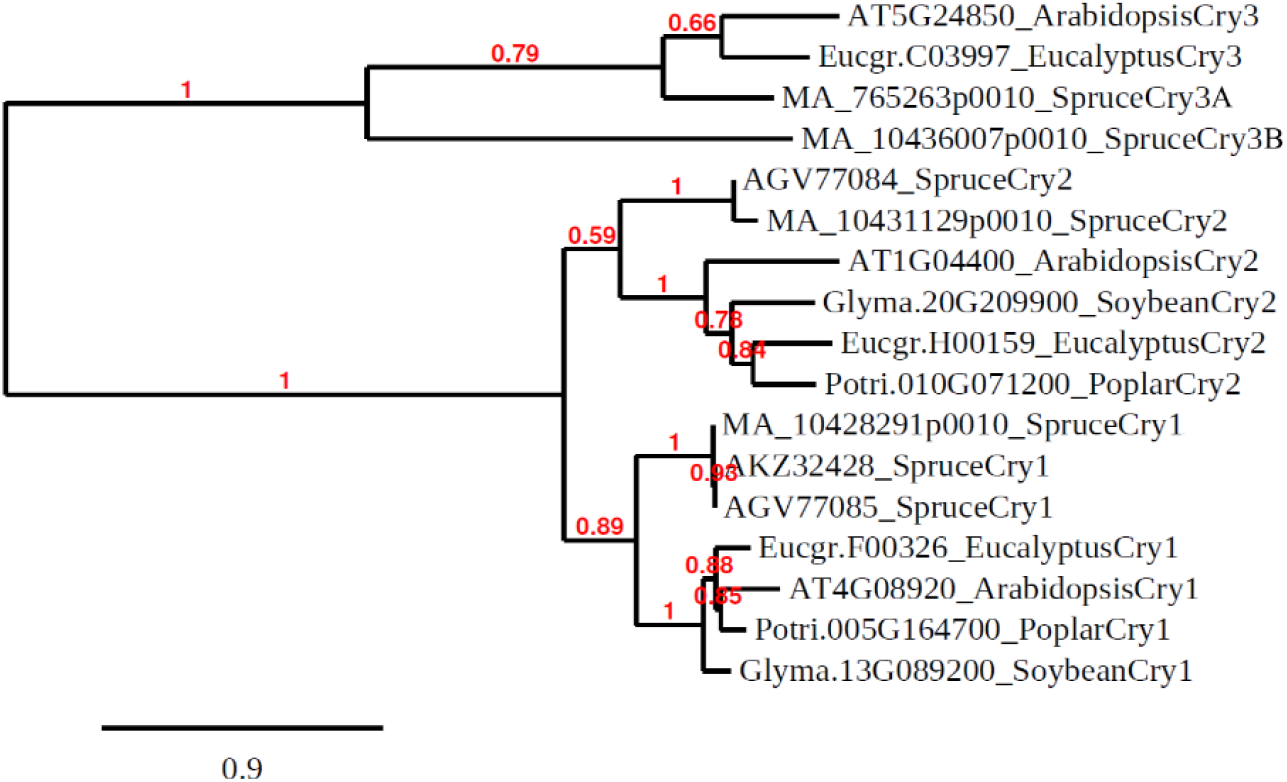
Phylogenetic tree of cryptochromes in Norway spruce

### Detection of SNPs in PHYs and CRYs

SNPs were detected in the coding regions of *PHYO, PHYP2, CRY1* and *CRY2*, whereas no SNPs were found in *PHYN, PHYP1* and *CRY3*. Thus, *PHYN, PHYP1* and *CRY3* serve as the control genes (photoreceptors) that did not show any polymorphism in their respective coding regions. One missense and one synonymous mutation, both with cline, were detected in *PHYO*. In *PHYP2*, two missense were found with cline, while two missense and two synonymous mutations were detected without cline. Regarding *CRY*, one missense with cline was detected in *CRY1*, while in *CRY2*, one missense with cline and one synonymous without cline were found (Table 1). The mutations without cline serve as additional controls confirming that only specific SNPs display the cline.

**Table 1.**
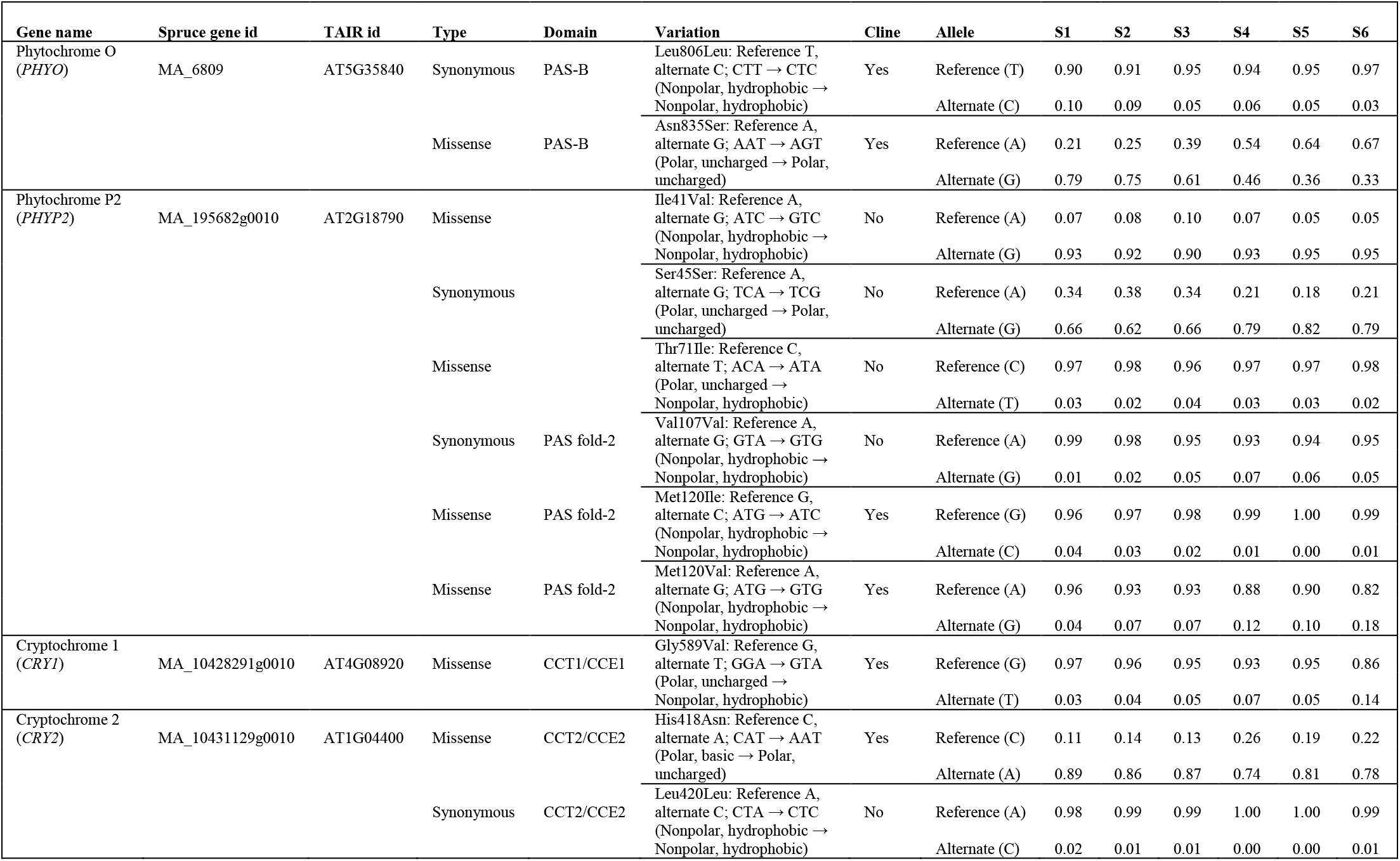
Phytochrome and cryptochrome SNPs in Norway spruce

Details of the position and type of SNP along with sequence information and allele frequencies for all four genes are represented in Table 1. The current work was focused on the missense SNPs in the coding regions, as these polymorphisms cause a change in the protein sequence, which may contribute to modify the protein conformation, leading to an alteration in their mode of action. The five missense SNPs with a cline in their allele frequencies and genotype frequencies belong to the conserved domains of the respective photoreceptors with well-defined functions. Asn835Ser polymorphism was detected in the PAS-B domain of PHYO. PAS is a short form from the names of the proteins in which imperfect repeat sequences were first recognized: *Drosophila* period clock protein (**P**ER), vertebrate aryl hydrocarbon receptor nuclear translocator (**A**RNT) and *Drosophila* single-minded protein (**S**IM) (Nambu et al. 1991). Met120Ile and Met120Val polymorphisms in the PHYP2 belong to the PAS fold-2 domain, while Gly589Val and His418Asn from CRY1 and CRY2 belong to the CRY1 C-terminal domain (CCT1) or CRY1 C-terminal extension (CCE1) and CRY2 C-terminal domain (CCT2) or CRY2 C-terminal extension (CCE2) respectively.

Three missense polymorphisms in the photoreceptors with cline resulted in alteration of the amino acid with similar chemical properties (Table 1). Asn835Ser (PHYO) involves a change in the amino acid with a similar chemical property; polar, uncharged Asn gets altered to Ser which is also polar and uncharged. Similarly, Met120Ile and Met120Val in PHYP2 resulted in a change of an amino acid that is nonpolar and hydrophobic to another amino acid with the same chemical property. Two missense SNPs were detected that involved the alteration of an amino acid to another amino acid with a different chemical property. Gly589Val (CRY1) resulted in the alteration of a polar, uncharged amino acid to a nonpolar, hydrophobic and His418Asn (CRY2) resulted in a change from polar, basic amino acid to a polar, uncharged one. The alteration of an amino acid to another amino acid with a different chemical property would modify the mode of action of the protein largely as compared to the alteration of the amino acid to another amino acid with a similar chemical property.

### Latitudinal clines in the allele and genotype frequencies of detected SNPs

The missense SNP in *PHYO* – Asn835Ser, displayed a statistically significant cline in the allele frequencies and genotype frequencies in the Norway spruce populations from different latitudes across Sweden (Figure 2) which is the steepest cline among the other missense SNPs in *PHYP2, CRY1* and *CRY2* that showed moderate clines (Table 1, Supplementary Information, Table S2, Table S3, Figures S12-S14). This is also evident from the highest *F*_*ST*_ value along with higher and less dispersed R^2^ values of *PHYO* as compared to the other photoreceptors (Figure 3). The pair-wise *F*_*ST*_ estimates for six populations of Norway spruce across Sweden (Table 2) including seven missense and four synonymous SNPs from four photoreceptors show that the *F*_*ST*_ values are low and show different levels of dispersion towards the north, which can be explained by an increase in differentiation within the populations with pair-wise geographic distance. A low *F*_*ST*_ value suggests low population genetic differentiation. The missense SNP loci from *PHYP2* and *CRY2* were found to be in HWE while only the southernmost population with reference to *CRY1* missense loci was in HWE (Supplementary Information Table S4). In the case of *PHYO*, only the southernmost and northernmost populations were found to be in HWE. This suggests that the respective loci in HWE will remain constant without undergoing an evolution in those populations from one generation to the next in absence of any external disturbing factors, while those from *PHYO* and *CRY1* will continue to evolve.

**Figure 3.**
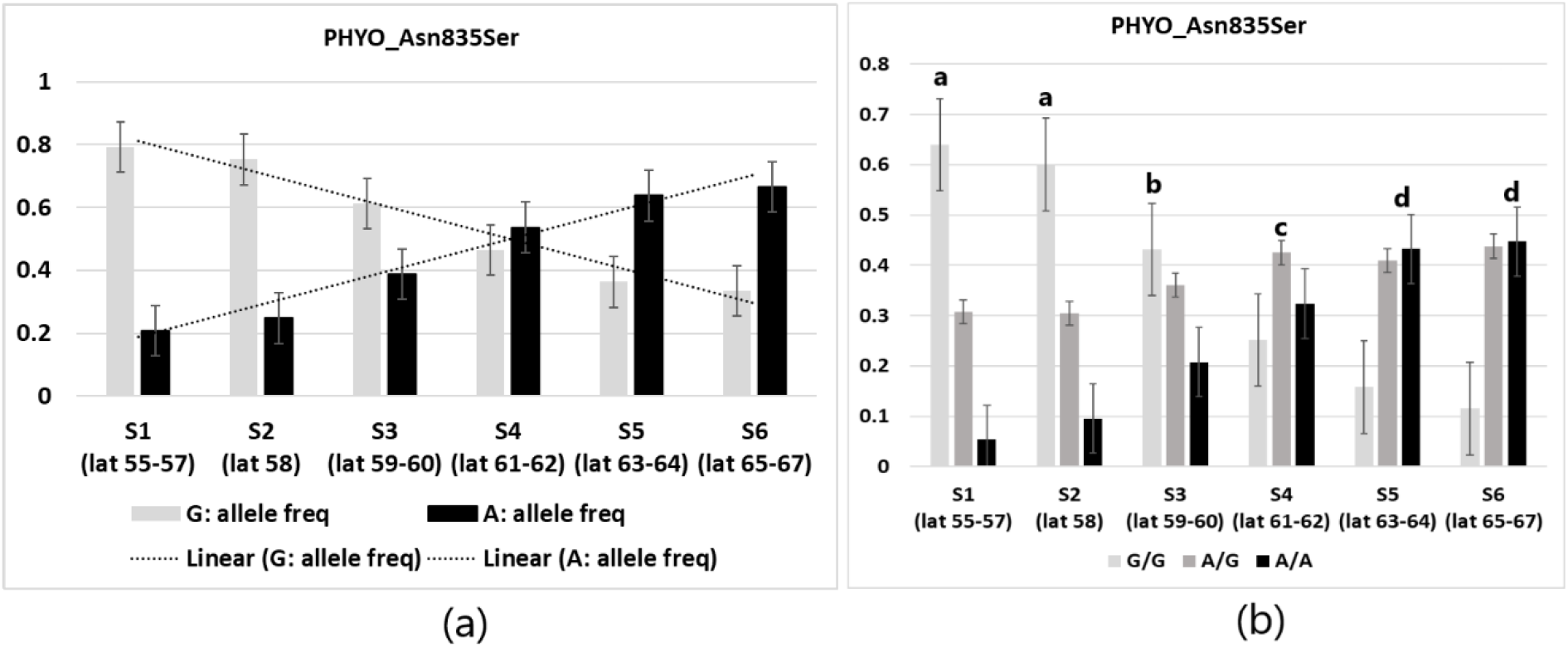
Cline regarding variation in allele and genotype frequencies of SNPs in the *PHYO* gene in Norway spruce populations across Sweden. **(a)** allele frequencies of Asn835Ser. **(b)** genotype frequencies of Asn835Ser, Tukey’s *post-hoc* categorization is indicated above the bars.

**Table 2.** Pairwise *F*_*ST*_ estimates for six populations of Norway spruce across Sweden involved in the analysis for detection of latitudinal clines in allele frequencies of the SNPs of phytochromes and cryptochromes

| Population | S2 | S3 | S4 | S5 | S6 |
| --- | --- | --- | --- | --- | --- |
| S1 | 0.00 | 0.005661 | 0.006079 | n.a. | 0.008693 |
| S2 |  | 0.00221 | 0.002378 | n.a. | 0.003764 |
| S3 |  |  | 0.00 | n.a. | 0.00 |
| S4 |  |  |  | n.a. | 0.00039 |
| S5 |  |  |  |  | n.a. |

## Discussion

From the phylogenetic analysis (Figure 1), the alignments (Supplementary Information, Figures S4-S7) and based on previous investigations in phytochromes (Mathews 2010; Schneider-Poetsch et al. 1998), it can be proposed that PHYO and PHYP from conifers are the equivalents of PHYA/PHYC and PHYB in angiosperms, respectively. The alignments show that the different domains in PHYs are well conserved in conifers. The PAS fold-2 domain of PHY is located in the N-terminal region that acts as a light sensing region, while PAS-A and PAS-B domains belong to the C-terminal region which is the regulatory region that is essential for dimerization, nuclear translocation and for modulating phytochrome signaling (Montgomery and Lagarias 2002; Rockwell et al. 2006). As the PAS domains are critical for light sensing and signaling, mutations in them lead to impaired/altered response to R/FR light (Neff et al. 2000; Paul and Kurana 2008). For example, missense mutations in the PAS fold-2 domain of PHYA from *Arabidopsis* showed impaired responses to R/FR light (Yanovsky et al. 2002; Wang et al. 2011) and Ile143Leu polymorphism in PAS fold-2 domain of PHYB from *Arabidopsis* was associated with variation in R light response (Filiault et al. 2008). Likewise, the CCT domain from *CRY* is involved in the blue light signal output (Yang et al. 2000; Wang et al. 2015). In the current study, missense SNPs detected in the coding regions of the respective domains may have an impact on the structure of the proteins altering their properties that may affect the mode of their action. The cline associated with these SNPs may be a probable explanation for the differential response to light quality of the Norway spruce populations from latitudes across Sweden that are exposed to variable light received at the respective latitudes (northern latitudes receive longer daily exposure to FR-enriched light as compared to the southern latitudes) (Clapham et al. 1998; Mølmann et al. 2006; Ranade and García-Gil 2021). The mutations causing a change of an amino acid to another amino acid with altered chemical property, e.g. Thr71Ile (PHYP2, Table 1), will alter the protein function largely, modifying its sensitivity to light quality. However, further investigation is required to validate the mechanism of the action of the altered PHY/CRY under variable light quality in Norway spruce, although this work provides a concrete model for the basis of latitudinal variation regarding differential response to light quality in Norway spruce populations, as photoreceptors are the core regulators of the light signaling pathway.

Nucleotide diversity in the photoreceptors was described by earlier investigations in Norway spruce. Although a significant excess of diversity was reported in photoreceptor genes such as *PHY* and *CRY* in Norway spruce previously, this study did not report any specific cline associated with the variation (Kallman et al. 2014). Latitudinal cline in the SNPs of a few genes that were differentially regulated under shade in different Norway spruce populations was reported earlier (Ranade and García-Gil 2021). Low *F*_*ST*_ values were reported by the earlier study (Ranade and García-Gil 2021), suggesting low population genetic differentiation, similar to the current analysis. Neither *PHY* nor *CRY* was found to be differentially regulated in Norway spruce in response to shade in the two populations from different latitudes (Ranade and García-Gil 2021). Norway spruce is well known for displaying a latitudinal cline in response to FR light (Clapham et al. 1998; Mølmann et al. 2006; Ranade and García-Gil 2021). The cline in the missense mutations in *PHY*s, especially the cline detected in *PHYO* (Asn835Ser) which is the steepest cline among all SNPs included in this analysis shows a strong correlation with the cline regarding the response to FR-enriched light reported earlier (Clapham et al. 1998; Ranade and García-Gil 2021) (Figure 5, Supplementary Information, Figure S2). A missense mutation causes a change in protein conformation altering its function. Therefore, although the *PHY*s were not differentially regulated, we propose that a change in their protein conformation due to the sequence polymorphism may affect its mode of action. This can be accounted for the difference in the requirement of FR in the Norway spruce populations across Sweden that follows a latitudinal gradient, which is concurrent with the cline in their SNPs.

**Figure 4.**
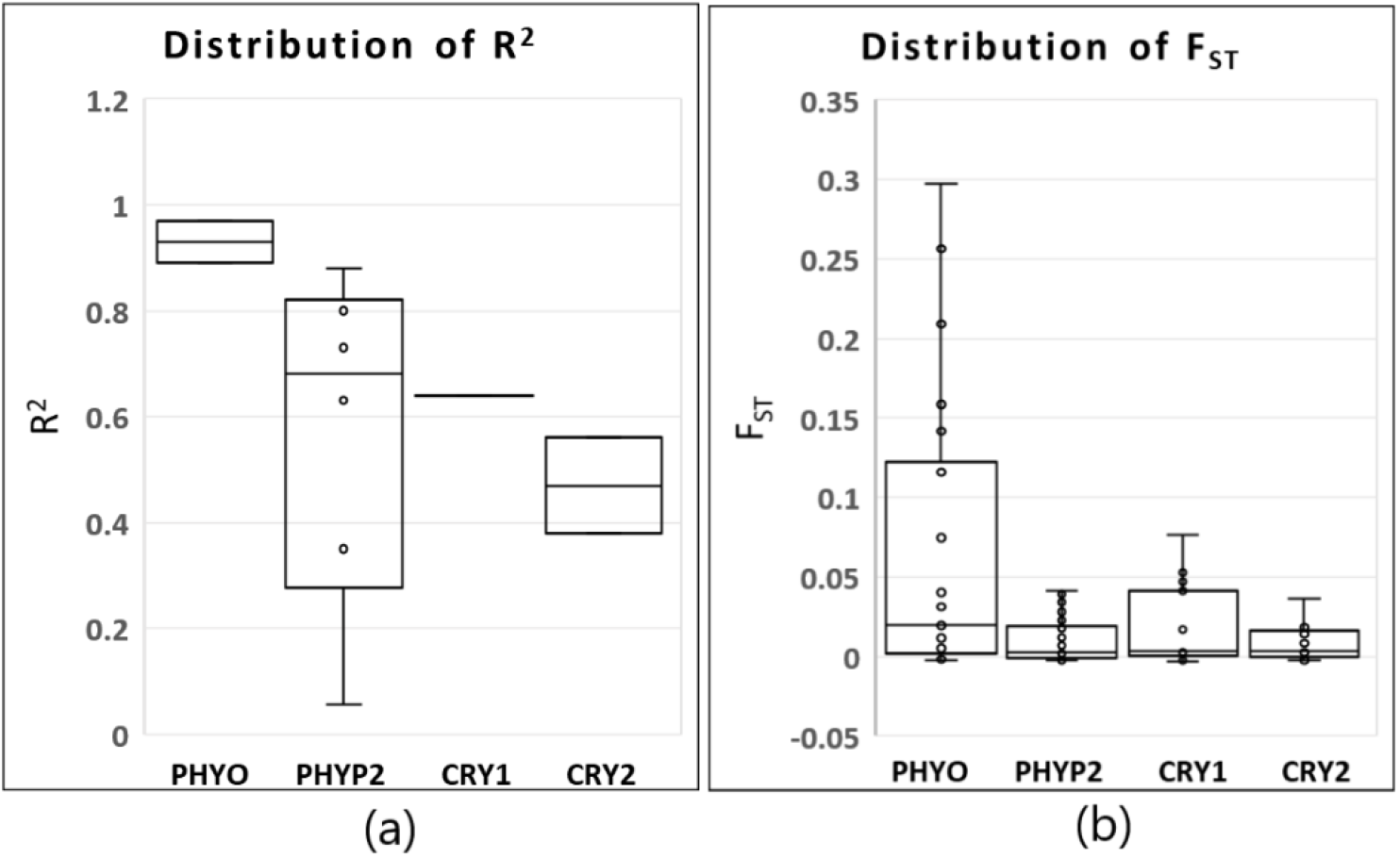
R^2^ and *F*_*ST*_ were calculated considering the missense and the synonymous SNPs detected in the particular candidate gene. (a) The distribution of R^2^ with reference to allele frequencies across the four photoreceptors showing clines in allele frequencies in the Norway spruce populations in Sweden. (b) The distribution of *F*_*ST*_ across the four photoreceptors showing clines in allele frequencies in the Norway spruce populations in Sweden.

**Figure 5.**
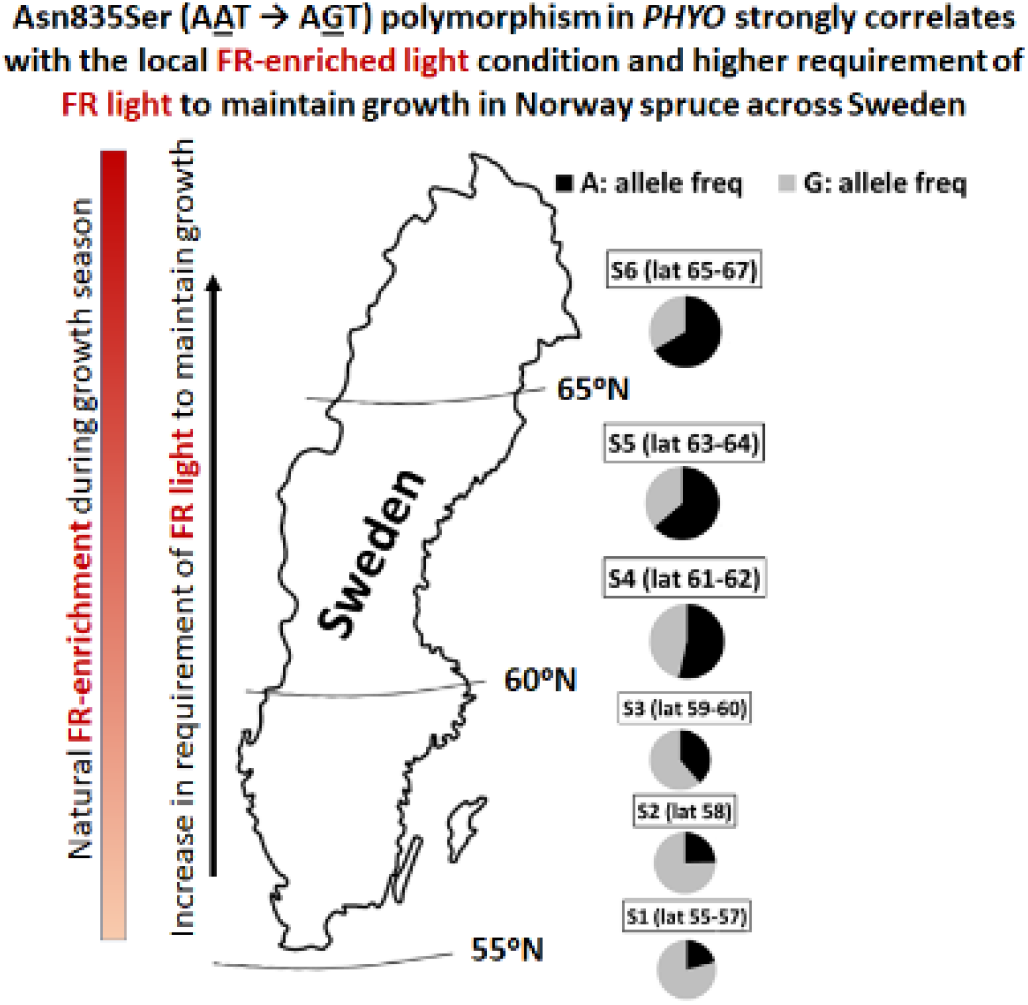
Cline regarding the requirement of FR light strongly correlates with the cline in Asn835Ser polymorphism in *PHYO* in the Norway spruce populations in Sweden. The pie charts in the figure represent the allele frequency of the polymorphism in the six Norway spruce populations included in the study.

From the current phylogenetic analysis and earlier reports (Mathews 2010; Schneider-Poetsch et al. 1998), it can be proposed that PHYO and PHYP from Norway spruce correspond to PHYA/PHYC and PHYB in angiosperms respectively. Duplication of *PHYA* resulted in the formation of *PHYC* and this duplication appears to have occurred before the diversification of angiosperms (Mathews and Sharrock 1997). Natural variation in *PHYs* associated with sensitivity to R/FR light has been identified in *Arabidopsis*, e.g. polymorphism in *PHYA* (Maloof et al. 2001) associated with FR and polymorphism in *PHYB* associated with R (Filiault et al. 2008). This further supports the current finding of natural variation in *PHYs* being linked with the earlier reported cline to the requirement for FR and clinal variation in response to low R:FR in Norway spruce as discussed before. PHYA and PHYB appear to have complementary functions in the processes related to seedling development and flowering (Reed et al. 1994). Clinal variation in *PHYB2* in *Populus tremula* was associated with growth cessation and bud-set (Ingvarsson et al. 2006), while control of growth cessation and bud-set by *PHYA* was demonstrated in hybrid aspen (Olsen et al. 1997). *Arabidopsis* mutants in CRY1/CRY2 are linked to differential response to photoperiod. A *CRY1* point mutation leads to early flowering under short day conditions and is hypersensitive to blue, R and FR light, in hypocotyl growth inhibition in *Arabidopsis* (Exner et al. 2010). Likewise, a variation in *CRY2* showed a difference in flowering response to photoperiod (El-Assal et al. 2001; Olsen et al. 2004). Particularly in *Picea*, genes harboring SNP associations with bud-set were found to be similar to *PHYA* and *CRY* in *Picea sitchensis* (Holliday et al. 2010). The latitudinal clines in response to light quality and the processes related to the response to photoperiods such as growth-cessation and bud-set/bud-burst have been described in Norway spruce populations across Sweden (Clapham et al. 1998; Sogaard et al. 2008), yet their association with SNPs in photoreceptors is not mentioned. The clines in the allele/genotype frequencies of the variations in photoreceptors observed in the current analysis may be the probable explanation for the clines related to growth-cessation and bud-set in Norway spruce reported by earlier investigations as these show a strong correlation with each other. Phytochromes function as thermosensors in plants, apart from sensing the light wavelengths (Hayes et al. 2021; Jung et al. 2016). There is a year-round sharp decrease in the temperatures across the latitudes towards the north. The polymorphisms in the phytochromes reported in this work can be also correlated with the temperature gradient prevalent across the latitudes, although this remains a hypothesis, which needs further validation.

PHYO from conifers is the equivalent of PHYA/PHYC in angiosperms as discussed in the earlier part of this article. PHYC perceives R/FR light and plays a major role in photomorphogenesis (Li et al. 2019) and is essential in photoperiod depended flowering, particularly under long day photoperiod (Chen et al. 2014; Woods et al. 2014). Natural variation in *PHYC* is associated with variation in flowering and growth responses in angiosperms (Balasubramanian et al. 2006; Saidou et al. 2009). PHYA mediates FR light promotion of flowering with modes of action similar to CRY2; both show a diurnal rhythm in short day plants acting as day sensors (Mockler et al. 2003). *Flowering locus T* (*FT*) plays a central role in the induction of flowering that is regulated by photoreceptors (Mockler et al. 2003; Bohlenius et al. 2006) and the photoreceptors in turn perceive and respond to light wavelength. An *FT* gene was found to be involved in the photoperiodic control of bud-set in a tree species – poplar (Bohlenius et al. 2006). A homolog of *FT* shows a significant correlation between its expression and bud-set or growth cessation in Norway spruce (Gyllenstrand et al. 2007; Karlgren et al. 2013). In addition, clinal variation was observed in *FT* expression levels that increased with latitude in Norway spruce (Chen et al. 2012). In this context, the cline with the missense polymorphism in *PHYO* in Norway spruce can be correlated with the latitudinal gradient in the expression levels of the *FT* gene along with the cline in response to variation in FR light, which needs further molecular validation.

## Conclusions

Light signaling regulates plant development throughout the plant life cycle. Light responses are mediated by the photoreceptors that play a central role. The cline in the SNPs in the *PHYs* detected in the current work strongly correlates with the latitudinal cline in response to R:FR ratio, cline regarding the higher requirement of FR and the cline associated with growth rhythms (bud-set/bud-burst) reported in earlier investigations in Norway spruce. The presence of a higher amount of FR in the northern latitudes during the growth seasons supports the adaptation of higher requirements of FR in the Norway spruce populations that thrive in these latitudes. However, further studies are required to confirm the molecular basis of the phenomenon. This is the first report so far in conifers, where precise and statistically significant clines were observed in the SNPs in *PHY*s that correlate strongly with response to light. We propose that these variations represent signs of local adaptation to light quality in Norway spruce. The knowledge gained by this analysis could be applied to designing strategies in breeding programs for Norway spruce.

## Abbreviations

CCT/CCE: *CRY* C-terminal domain/*CRY* C-terminal extension
CRY: Cryptochrome
FR: Far-red light
PAS: *Drosophila* period clock protein (**<u>P</u>**ER), vertebrate aryl hydrocarbon receptor nuclear translocator (**<u>A</u>**RNT) and *Drosophila* single-minded protein (**<u>S</u>**IM)
PHY: Phytochrome
R: Red light
SNP: Single nucleotide polymorphism

## Author contributions

Both authors contributed with experimental design, data analysis and interpretation, and manuscript writing. Both authors read and approved the manuscript.

## Acknowledgments

We acknowledge the support from Science for Life Laboratory (SciLifeLab), the Knut and Alice Wallenberg Foundation, the National Genomics Infrastructure funded by the Swedish Research Council, and Uppsala Multidisciplinary Centre for Advanced Computational Science for assistance with massively parallel sequencing and access to the UPPMAX computational infrastructure. We acknowledge Swedish Foundation for Strategic Research (SSF) for their support regarding the exome capture project. We also acknowledge Swedish Research Council (VR) and Swedish Governmental Agency for Innovation Systems (VINNOVA) for their support.

